# Improvement of the potency of a N1-methylpseudouridine-modified self-amplifying RNA through mutations in the RNA-dependent-RNA-polymerase

**DOI:** 10.1101/2024.06.14.599087

**Authors:** Verónica Quintana, Josefina Caillava, Laura A. Byk, Juan A. Mondotte, Leandro Battini, Prutha Tarte, Marcelo M. Samsa, Claudia V. Filomatori, Diego E. Alvarez

**Author notes:** These authors contributed equally (Verónica Quintana, Josefina Caillava).

## Abstract

RNA vaccines are sensed as non-self molecules by the innate immune system, and balancing control of the immune activation and vaccine safety and efficacy has remained a challenge, especially for self-amplifying RNAs (saRNAs). Incorporation of modified nucleotides has been widely used to temper immune activation of RNA vaccines. However, it was previously reported that incorporation of modified nucleotides to saRNAs impeded antigen expression. Here, we used a reporter replicon of the attenuated TC-83 strain of Venezuelan equine encephalitis virus (VEEV) to investigate the impact of modified nucleotide incorporation on the replication capacity of the saRNA in transfected cells. ψ and ψ-modified molecules showed a profound defect in RNA synthesis compared to unmodified saRNA. Interestingly, the levels of RNA synthesis of m^5^C-modified RNAs were similar to unmodified molecules, positioning m^5^C as a promising candidate for saRNA modification. To overcome the impact of ψ or m^1^ψ-modified nucleotide incorporation in RNA synthesis, we explored two alternative approaches: engineering the UTR sequences and tuning polymerase fidelity. Our results uncover a previously unappreciated link between polymerase fidelity and saRNA amplification. Overall, we provide new insights for the design of saRNAs with high levels of heterologous protein expression and potential vaccine applications.

## INTRODUCTION

Vaccines and immunotherapies based on RNA molecules rely on the biological role of RNA as a messenger (mRNA) for protein translation by the host cell to achieve native payload expression including post-translational modifications, assembly of multimeric protein complexes, and appropriate trafficking to subcellular locations. The rapid development and simple production process by *in vitro* transcription, the lack of risk of integration into the host DNA genome, and the proven effectiveness compared to other vector-based platforms and inactivated viral vaccines are major advantages of RNA-based platforms for vaccine development [1–3]. However, mRNA vaccine technology faces RNA instability, potent activation of the innate immune response that limits RNA translation, and high dose requirements often resulting in higher rates of side effects compared to other vaccine platforms. Different strategies aim at increasing the yield of antigen expression upon delivery of RNA molecules by controlling immune activation or improving translation [1]. First, incorporation of type 1 or 2 cap during the *in vitro* transcription or enzymatically after RNA synthesis mimics endogenous mRNA molecules, limiting the intrinsic immune response. Second, the 5’ and 3’ untranslated regions (UTRs) can be optimized to enhance translational efficiency and control immune response. Third, incorporation of modified nucleotide analogues including 5-methylcytidine (m^5^C), N6-methyladenosine (m^6^A), 5-methyluridine (m^5^U), 2-thiouridine (s^2^U) or pseudouridine (ψ) is a commonly used strategy aimed at reducing the activation of the immune response in transfected cells [4]. In addition, ψ and N1-methylpseudouridine (m^1^ψ) increase the translational capacity of modified mRNAs [5]. Other strategies such as codon optimization of the open reading frame (ORF) encoding the protein of interest or increasing the length of the poly(A) tail, are also applied with disparate outcomes. Finally, vaccine design based on self-amplifying RNAs (saRNAs) provides means to lower dose requirements due to the saRNAs’ capacity to replicate in the cell cytoplasm, generating multiple copies of translatable RNAs and assuring high yields of heterologous proteins [6,7].

saRNAs are derived from the genomes of positive-strand RNA viruses. The most usual design follows the organization of alphavirus genomes, with viral 5’ and 3’UTRs at the ends of the saRNA molecules [8]. A first ORF encodes for the replicase complex consisting of non-structural proteins 1-4 (nsP1-4), and a second ORF, under the control of a subgenomic promoter, encodes the vaccine antigen of interest in place of the original viral structural proteins. Upon delivery into the cell cytoplasm, saRNA molecules are first translated to produce the nsP1-4 proteins that assemble into the replicase complex. The newly assembled replicase complex uses the originally delivered molecule as a template to transcribe a complementary RNA copy that in turn serves as the template for the transcription of full-length and subgenomic RNAs. Therefore, the protein of interest is only translated from the subgenomic RNA once the input RNA is transcribed and amplification is achieved within a few hours in the following rounds of replication.

Innate immune activation is a major issue in saRNA technology. In addition to the high immunogenicity of the synthetic molecule delivered as a vaccine, a cell response is elicited against double stranded RNA (dsRNA), replication intermediates and viral antigens [9]. Given that expression of the antigen requires the initial translation of replicase components, the immune response mounted upon RNA delivery into the host cell determines the yield of antigen expression. Noteworthy, although saRNAs are attractive systems for the delivery of heterologous antigens, strategies to reduce the immunogenicity of mRNA vaccines, such as through the incorporation of modified nucleotides or sequence engineering, are not accepted by saRNA molecules [10,11]. As a matter of fact, antigen expression following intramuscular injection of ψ-modified saRNA, derived from Venezuelan equine encephalitis Virus (VEEV), encoding recombinant Zika virus monoclonal antibody was undetectable in mice [10]. The underlying impact of modified nucleotide incorporation leading to the loss of function of saRNAs remains largely unexplored both *in vitro* and *in vivo*. Altogether, unlike the two licensed COVID-19 mRNA vaccines or majority of vaccine candidates in clinical trials that use modified nucleotides analogues to limit innate immune activation, saRNA vaccine candidates that have advanced into clinical trials use unmodified RNAs [8]. The use of sequence optimization to modulate immune response is also constrained for saRNAs due to the strict sequence and structure requirements of cis-acting replication elements that map both to viral UTRs and coding sequences [12].

Optimization of replicase components remains an attractive alternative to improve payload expression from saRNA. Following nsP1-4 translation, the alphavirus replicase assembles into specialized membrane compartments of the infected cells [13]. Nsp1-4 is expressed as a polyprotein, and initial cleavage by nsP2 at the junction between nsP3 and nsP4 results in the formation of the early replication complex that copies a complementary negative-strand of the full-length RNA. The late replication complex forms after further processing of nsP1-3 into individual proteins: nsP1 mediates anchoring of the complex to the plasma membrane and is required for capping of viral RNAs, nsP2 functions as an RNA helicase and viral protease, nsP3 serves as a scaffold for the recruitment of host factors, and nsP4 is the RNA-dependent-RNA-polymerase (RdRp). *In vitro* evolution of VEEV replicons has prompted the identification of defined mutations in nsp2 and nsp3 that enhance heterologous protein expression *in vitro* and *in vivo* [14].

Here, we investigated the impact of incorporating modified nucleotides into a saRNA molecule based on the TC-83 attenuated strain of VEEV. We observed that modified RNAs are translated efficiently but have impaired levels of RNA synthesis in cell culture. Mutations designed to alleviate the effect of modified nucleotide incorporation in the UTRs were not able to rescue replication levels. Alternatively, with the aim of engineering a viral replicase capable of tolerating modified nucleotides, we introduced substitutions in the nsp4 RdRp that have been reported to alter fidelity [15,16]. Importantly, we found that replicons carrying mutations associated with a high-fidelity polymerase phenotype were able to replicate m^1^ψ-modified saRNAs at higher levels and displayed higher infectivity than wild type replicons. Altogether, our data introduce a new notion linking polymerase fidelity to its ability to accept modified RNAs as templates for RNA synthesis.

## RESULTS

### Assessment of translation and replication ability of VEEV-based saRNAs incorporating modified nucleotides

To gain insight into the impact of incorporating modified nucleotides on saRNA translation and replication, we designed a reporter VEEV-based replicon carrying the Renilla luciferase gene inserted into the hypervariable domain of nsp3 in the first ORF, and the firefly luciferase gene followed by the 2A autoprotease of foot and mouth disease virus (FMDV) and GFP genes in place of the viral structural proteins in the second ORF (Figure 1A). In this construct, Renilla luciferase is translated from the input RNA upon transfection once the saRNA is delivered inside the cell. In turn, firefly luciferase is translated from the subgenomic RNA that is produced by the viral replicase after an initial round of translation and RNA synthesis. In addition to the wild type replicon, we constructed a mutant replicon, bearing a mutation in the nsp4 RdRp catalytic site that turns the polymerase inactive.

**Figure 1.**
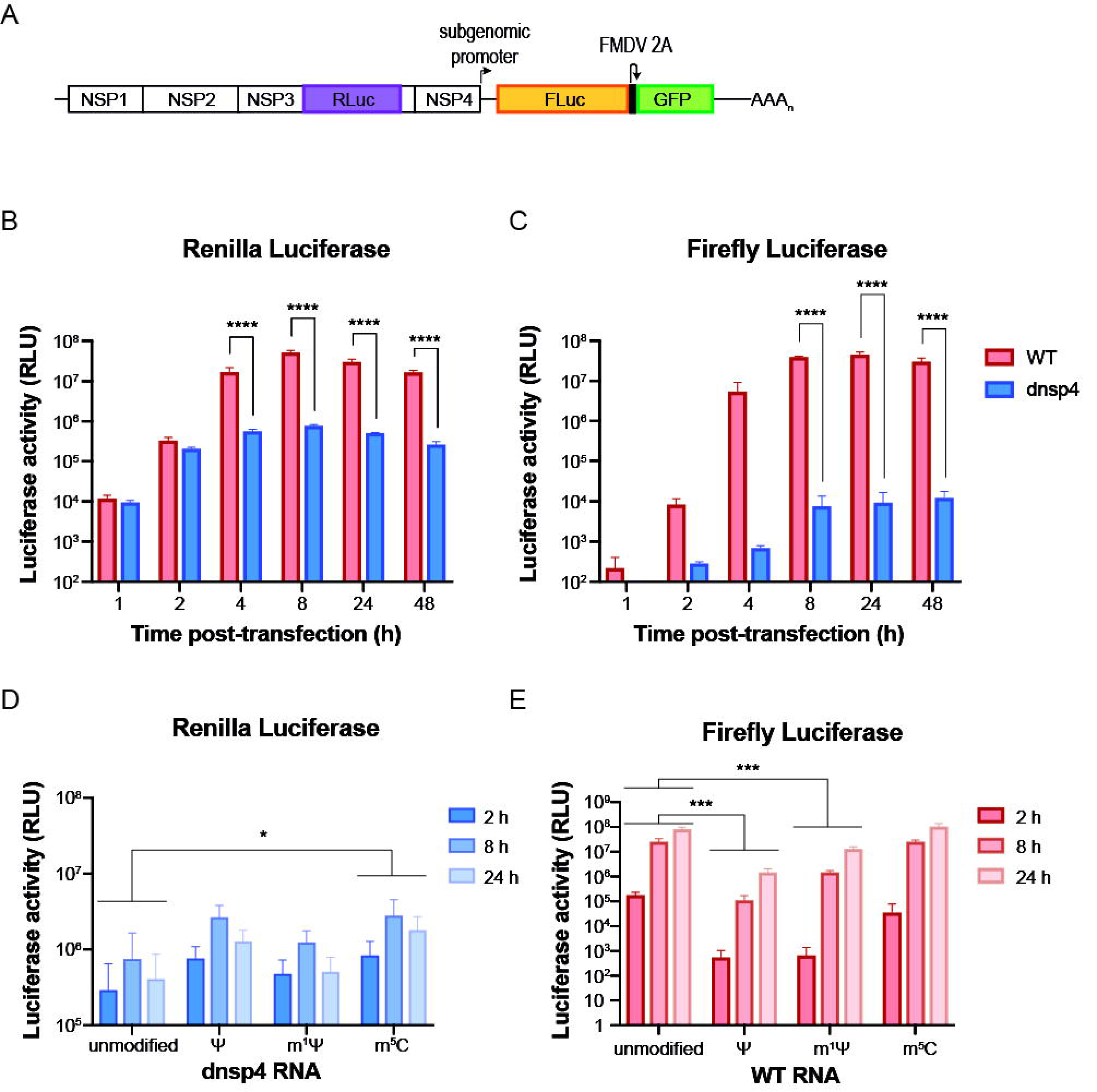
Uridine analogs impair the self-amplifying ability of a VEEV-based replicon RNA. (A) Schematic representation of a VEEV TC-83 -based reporter replicon. Renilla luciferase sequence (RLuc) was inserted in the hypervariable region of nsp3. Firefly luciferase (FLuc) and GFP sequences separated by FMDV 2A autoprotease cleavage sequence were inserted in place of the viral structural proteins under the control of the subgenomic promoter in the second ORF. (B) and (C) Bar graph showing Renilla (B) and firefly (C) luciferase activity as a function of time after transfection of BHK-21 with 80ng RNA of VEEV replicon (WT, red bars) or catalytically inactive nsP4 mutant (dnsp4, blue bars). (D) and (E) Bar graph displaying Renilla luciferase activity of dnsp4 construct (D) and firelfy luciferase activity of WT (E) at 2-, 8- and 24-hours post-transfection of VEEV replicon RNA transcribed in vitro with unmodified nucleotides or with ψ, mψ or 5mC. Two independent experiments with two replicates per condition were analyzed. Statistical analysis was performed using two-way ANOVA, followed by Sidak’s (B and C) test to compare means at each time point or Dunnett’s (D and E) test to compare the main effect of modified nucleotides vs. unmodified. ns p> 0.05; *p ≤ 0.05; **p ≤ 0.01; ***p ≤ 0.001; ****p ≤ 0.001. Error bars indicate the standard deviation.

First, the wild type (WT) and catalytically inactive nsp4 mutant (dnsp4) saRNAs synthesized *in vitro* were transfected into BHK-21 cells to characterize the expression kinetics of the luciferase reporter genes (Figure 1B and C). Renilla luciferase activity was detected as early as one hour after transfection, displaying similar levels for the WT and the dnsp4 constructs. Therefore, early Renilla luciferase activity was used to estimate the translation of the input RNAs. At later time points, Renilla activity increased for both constructs and peaked at eight hours post-transfection. At four hours post-transfection, Renilla luciferase levels for the WT and the dnsp4 constructs were significantly different, reflecting genomic RNA amplification by an active replicase complex (Figure 1B). In turn, firefly luciferase activity of the WT replicon was detected at low levels above background at one hour and it was readily detectable two hours post-transfection (Figure 1C). Thereafter, firefly luciferase levels for the WT construct increased exponentially and reached a maximum eight hours post-transfection. Therefore, firefly luciferase activity was used to estimate RNA replication and transcription of subgenomic RNA. In contrast, firefly luciferase levels detected for the dnsp4 construct marginally increased throughout the time course of the experiment and remained 1000-fold below WT levels, suggesting that the mutated nsP4 may retain leaky RdRp activity (Figure 1C). Altogether, our reporter saRNA molecules allowed monitoring translation of the input RNA by means of Renilla luciferase activity at an early time point. Meanwhile, measurement of firefly luciferase activity for the WT replicon, after two hours post-transfection, reflects RNA synthesis. In turn, Renilla luciferase activity for the dnsp4 construct reflects translation and stability of the input RNA.

To investigate the impact of the incorporation of nucleotide analogs on saRNA amplification, unmodified or modified RNAs of the WT and the dnsp4 mutant replicons were produced. Modified RNAs were *in vitro* transcribed using 100% ψ (pseudouridine), 100% m^1^ψ (N1-methylpseudouridine) in place of uridine or 100% m^5^C (5-methylcytosine) in place of cytidine. We monitored Renilla luciferase levels at 2-, 8- and 24-hours post-transfection for the dnsp4 mutant to assess translation and stability of the input RNA (Figure 1D). ψ- and m^5^C-modified RNAs displayed a similar overall trend of increased luciferase levels compared to unmodified RNA throughout the time course, while incorporation of m^1^ψ yielded luciferase levels similar to the unmodified RNA (Figure 1D) [5,17]. Using firefly luciferase levels to assess RNA replication, we observed that incorporation of ψ and m^1^ψ resulted in a reduction of luciferase activity (Figure 1E). The effect was significant and showed a 100-fold decrease in luciferase levels as early as 2 hours post-transfection and throughout the time course, compared to the unmodified RNA. Remarkably, m^5^C-RNA showed luciferase levels comparable to the unmodified RNA (Figure 1E). These results indicate that ψ and m^5^C have a moderate effect favoring translation and stability of the input RNA, and uridine analogues, namely ψ and m^1^, impaired the replicase function.

It has been widely reported that incorporation of nucleotide analogues diminishes immune sensing of foreign RNAs [5,17]. Therefore, we hypothesized that modified RNAs may have an advantage over unmodified RNAs in the context of cells possessing intact antiviral sensing and interferon signaling pathways. To investigate this possibility, A549 cells were transfected with modified or unmodified WT or dnsp4 RNAs, and Renilla and firefly luciferase activities were measured at 2-, 8- and 24-hours post-transfection (Figure S1). Compared to BHK-21 cells, Renilla luciferase levels in A549 cells suggested a trend of apparent increased translation, not only for ψ- and m^5^C-modified RNAs but also for m^1^ - modified RNA, relative to unmodified RNAs (Figure S1A). Assessment of replication by means of firefly luciferase activity measurement yielded results similar to those observed in BHK-21 cells, where incorporation of ψ and m^1^ resulted in overall reduced replication levels, while m^5^C had no apparent effect on RNA replication (Figure S1B).

Altogether, we used saRNA reporter constructs to discriminate between translation and stability of input RNA from RNA synthesis. Similar to mRNAs, incorporation of modified nucleotides into the saRNA resulted in enhanced translation and stability of the input RNA. However, ψ- and m^1^ψ-modified RNAs had reduced self-amplifying ability. Unlike uridine analogues, m^5^C-modified RNA displayed levels of self-amplifying capacity comparable to unmodified RNA.

### Optimization of the UTR sequences

The 5’ and 3’ ends of plus-strand RNA virus genomes often control RNA translation and synthesis. In effect, sequences in the VEEV 5’UTR promote negative-strand RNA synthesis through interaction with the viral 3’UTR, and a 51-nt conserved sequence element (CSE), downstream to the translation start codon, stimulates RNA replication. In the 3’UTR, a 19-nt CSE and the poly(A) tail are required for negative-strand RNA synthesis [12]. Noteworthy, usage of modified nucleotides in IRESs (Internal ribosome entry sites) and riboswitches was reported to alter their functionality [18,19]. It is therefore possible that the incorporation of ψ and m^1^ψ into viral UTRs impaired their function as promoters for RNA synthesis.

In particular, the uridine residue at the second position of the 5’ end of the genome is strictly required for alphavirus replication [20]. Modified RNAs co-transcriptionally capped using the trinucleotide analogue CleanCap result in the incorporation of an unmodified uridine at position 2 (U2) of the viral genome. In contrast, modified RNAs capped with a dinucleotide cap analogue carry a ψ in this position (ψ2) (Figure 2A). In previous studies in which alphavirus-derived modified saRNAs were reported to be non-functional *in vivo*, these RNAs were enzymatically capped after *in vitro* transcription with the dinucleotide analogue [10]. To assess whether the incorporation of ψ2 alters saRNAs’ functionality, we generated saRNAs in the context of our reporter construct carrying unmodified uridine or ψ in the second position by co-transcriptionally capping with Clean Cap or an ^m7^GpppA_m_ dinucleotide cap analog, respectively. To examine RNA translation and synthesis, unmodified or ψ-modified RNAs were transfected into BHK-21 cells. Renilla and firefly luciferase activities were measured at 1 and 24 hours after transfection, respectively (Figure 2B and C). One hour post-transfection, all four RNAs were efficiently translated and the impact of ψ incorporation was comparable for either of the capping strategies, resulting in a reduction of Renilla luciferase activity by about 3-fold (Figure 2B). Regardless of the capping analogue incorporated in the *in vitro* transcription, firefly luciferase activity at 24 hours indicated that ψ incorporation had a profound negative effect on saRNAs replication and led to a 100-to 200-fold decrease in luciferase levels for modified RNAs compared to the unmodified counterparts (Figure 2C). Altogether, the data suggest that replication of modified saRNA is impaired regardless of whether U or ψ is incorporated in the second position. Therefore, it is likely that the impact of ψ on saRNA replication is due to the incorporation of the ψ analogue across the RNA template.

**Figure 2.**
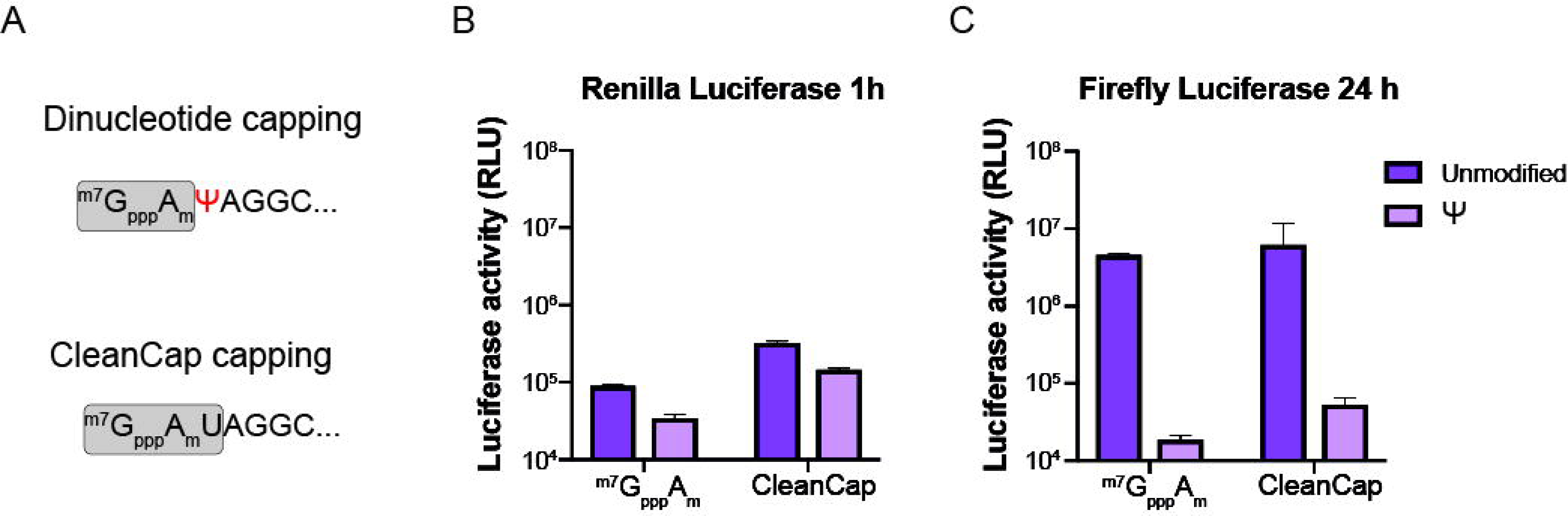
The strictly conserved uridine in the second position of VEEV 5’UTR tolerates. ψ **modification.** (A) Nucleotide sequence of the VEEV replicon 5’ end capped with a dinucleotide cap analog (top) or CleanCap (bottom). Grey boxes highlight cap structures. Positions incorporating ψ in the *in vitro* transcription are indicated in red. (B) and (C) Bar graphs of luciferase activity in BHK-21 cells transfected with 80 ng of unmodified or ψ-modified RNA cotranscriptionally capped with dinucleotide cap analog (m7GpppAm) or CleanCap. Renilla luciferase activity was measured 1h post-transfection (A) and firefly luciferase activity 24h post-transfection (B). One representative experiment out of two is shown.

The 5’UTR of VEEV folds into a stable stem-loop structure (5’SL), consisting of a small stem that exposes an 11 nt-long loop [21]. Changes in the stability or the sequence of the 5’SL have been shown to strongly affect virus replication. In fact, the attenuated TC-83 strain of VEEV, derived from the virulent Trinidad Donkey (TRD) virus, carries a mutation in the third position of the genome (G3A) that disrupts the stem base pairing, altering the structure of the 5’SL. As a result, genomic and subgenomic RNA levels are affected and their relative ratio is altered [21]. In addition to its impact on promoter activity, G3A results in an increased sensitivity to interferon (IFN) of TC-83 compared to the TRD strain [22]. The biological function of structured RNA elements often depends on their secondary structure and/or nucleotide sequences exposed in loops and bulges. Therefore, we reasoned that the 5’SL structure might be sensitive to the incorporation of nucleotide analogues, providing an explanation for the impaired replication of modified saRNAs. To address this possibility, we first designed a set of conservative mutations in the loop and in the region downstream of the 5’SL in the 5’UTR that preserved the 5’SL structure: uridines were replaced by adenosines, and cytidines by either adenosines or uridines (Figure 3A). RNA molecules from this set of mutants were *in vitro* transcribed with unmodified or modified nucleotides (m^1^ψ or m^5^C) and transfected into BHK-21 cells. No significant differences in Renilla luciferase levels were observed either for unmodified RNAs or for the modified RNAs of the wild type (5’UTR TC-83) and 5’UTR mutant constructs at 1-hour post-transfection (Figure 3B, left). The overall trend of Renilla luciferase expression for unmodified RNAs compared to their modified counterparts followed a similar pattern (Figure 3B, left). In all cases, luciferase levels were lower for m^1^ψ-RNAs, while levels for m^5^C-RNAs were similar for unmodified RNAs. This result suggests that mutations in the 5’SL did not confer an advantage for the translation of the input RNA, regardless of the presence of modified nucleotides. At 24 hours post-transfection, unmodified RNAs of the 5’UTR TC-83 and 5’UTR mutant constructs displayed similar levels of firefly luciferase activity (Figure 3B, right). Meanwhile, incorporation of m^1^ resulted in a 10-to 20-fold reduction in luciferase levels for the 5’UTR TC-83 and all of the 5’UTR mutant constructs, indicating that none of the mutations alleviated the effect of incorporating m^1^ into the saRNA. Similar to unmodified RNAs, mutations in m^5^C-RNAs had no impact on firefly luciferase levels. These results confirm that incorporation of m^1^ψ negatively affects viral replication, and its effect cannot be rescued by conservative mutations at uridine and cytosine positions that avoid modified analog incorporation but maintain the 5’SL structure.

**Figure 3.**
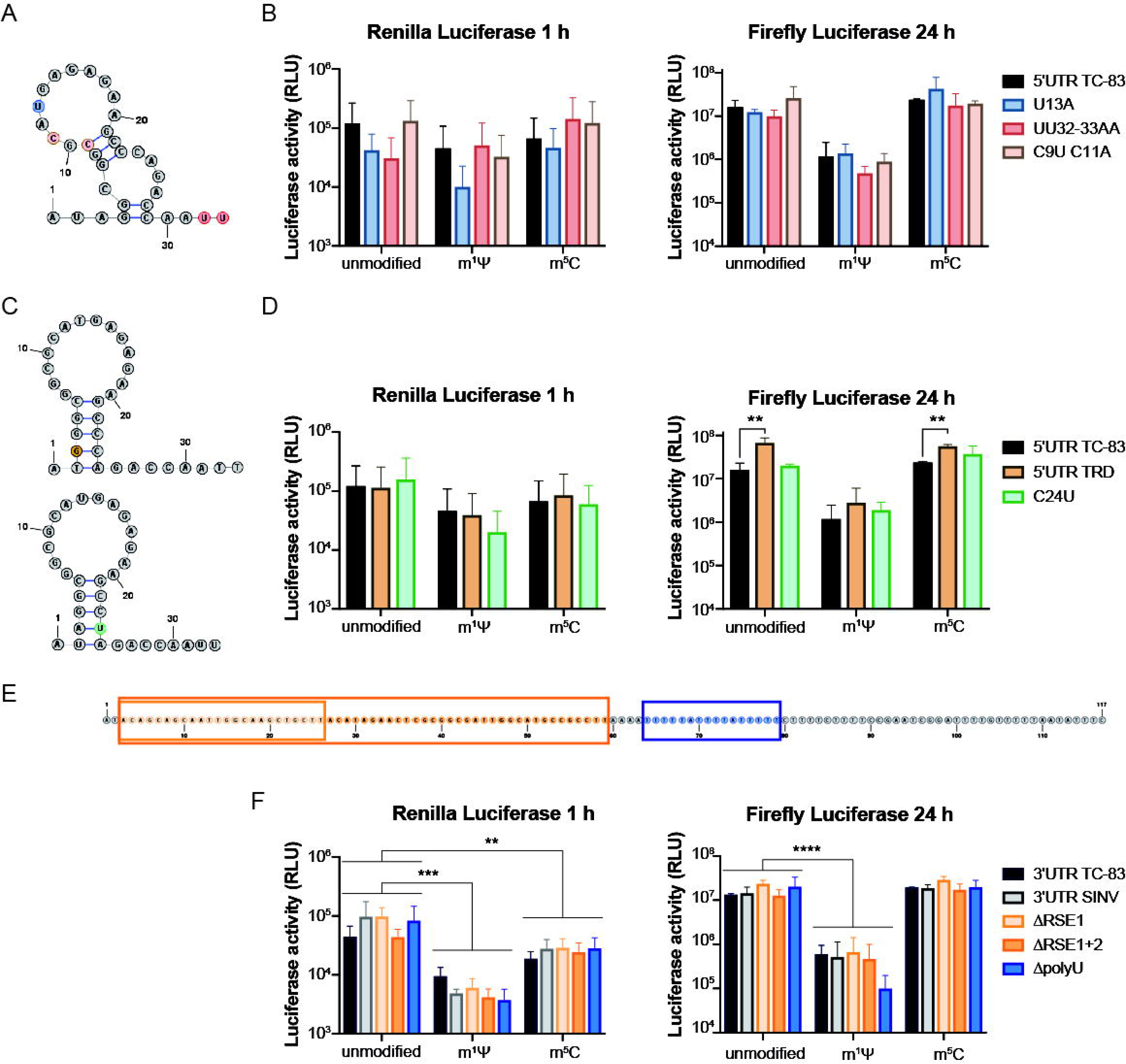
Designed 5’ and 3’UTR mutants do not alleviate the impact of nucleotide modifications on the self-amplifying ability of VEEV replicons. (A) Predicted secondary structure of TC-83 5’UTR. The indicated nucleotide changes were designed to preserve the overall secondary structure. (B), (D) and (F) Bar graphs showing luciferase activity of BHK-21 cells transfected with 80 ng of unmodified RNA, or m1ψ or 5mC-modified RNAs. Renilla luciferase activity was measured at 1h post-transfection (left) and firefly luciferase activity was measured at 24h post-transfection (right). (C) Predicted secondary structure of TRD 5’UTR (top) and C24U point-mutation (bottom). (E) Nucleotide sequence of TC-83 3’UTR. Boxes indicate the regions deleted in the designed mutants. Two independent experiments with two replicates per condition were analyzed. Statistical analysis was performed using two-way ANOVA, followed by Dunnett’s multiple comparisons test. Zig-zag lines in (D) compare the effect of mutations against the wild type within unmodified, m^1^ψ-, or m^5^C-modified RNAs. Statistics in (B) and (F) compare the main effect of modified nucleotides against unmodified. ns p> 0.05; **p ≤ 0.01; ***p ≤ 0.001; ****p ≤ 0.001. Error bars indicate the standard deviation.

Next, we tested the functionality of TRD VEEV stem loop in the TC-83 background (i) by replacing adenine in the third position by guanosine (A3G) to reconstitute the 5’UTR TRD sequence or (ii) by mimicking TRD stem loop structure mutating cytosine at position 24 in TC-83 5’UTR by uridine (C24U) (Figure 3C). In both mutants, the 5’UTR folds into a TRD-like 5’SL with comparable stability. According to previous studies, the TRD 5’SL is capable of counteracting the host antiviral response and maintaining full promoter functionality [22]. Renilla luciferase activity levels one hour after transfection of the 5’UTR TRD or the C24U mutant were similar to the parental 5’UTR TC-83 saRNA and followed the same trend described above after incorporation of m^1^ψ or m^5^C into the input RNA (Figure 3D, left). As expected, at 24 hours post-transfection, unmodified 5’UTR TRD displayed a significant 4-fold increase in firefly luciferase levels compared to the parental 5’UTR TC-83 (Figure 3D, right). Notably, an increase was also observed for m^1^ψ- or m^5^C-modified 5’UTR TRD RNAs. In turn, luciferase activity levels of unmodified and modified RNAs remained unaltered for the C24U mutation compared to the parental 5’ UTR TC-83. In conclusion, restoring the 5’UTR TRD sequence and structure conferred a replicative advantage to both unmodified and modified saRNAs. However, a 5’SL with an artificial nucleotide sequence predicted to reconstitute the TRD structure did not favor RNA replication.

Finally, we set out to optimize the 3’UTR sequences. To this end, we designed a set of four mutants. In the first mutant, we replaced the 3’UTR of TC-83 VEEV by the 3’UTR of Sindbis virus (SINV), which has been implicated in alphavirus RNA stabilization in mammalian cells [23]. In addition, VEEV replicase complex was shown to replicate in trans, a minimal template carrying SINV promoter sequences including the 5’ and 3’ UTRs [24]. The other three mutants contained deletions in the TC-83 3’UTR (Figure 3E). The 3’UTR of TC-83 contains two copies of a repeated sequence element (RSE), followed by a uridine-rich sequence tract. For alphaviruses that alternate between humans and mosquitoes, such as chikungunya and Eastern Equine Encephalitis, the RSEs have been found to be redundant or even unfavorable for viral replication *in vitro* in mammalian cells [12]. In turn, uridine content has been linked to the immune stimulating activity of mRNAs [25]. Based on these previous observations, we engineered deletions to remove one or two copies of the RSEs, or the entire poly(U) sequence (Figure 3E). Then, *in vitro* transcribed unmodified or modified saRNAs were transfected into BHK-21 cells. One hour after transfection, unmodified RNAs of the wild type (3’UTR TC-83) and 3’UTR mutant replicons displayed comparable levels of Renilla luciferase activity, indicating that the 3’UTR mutations did not affect the saRNA translation (Figure 3F, left). Incorporation of m^1^ψ resulted in an overall 10-fold decrease in Renilla luciferase levels for the wild type and the whole set of mutant replicons, while incorporation of m^5^C impacted on luciferase levels to a lesser extent. At 24 hours, 3’UTR TC-83 and 3’UTR mutant m^1^ψ-RNAs displayed 10-to 100-fold lower firefly luciferase levels than their unmodified counterparts, while m^5^C-RNAs showed similar levels of luciferase activity to the unmodified molecules (Figure 3F, right).

Because luciferase levels reflect the bulk expression of the payload in the transfected cells, we decided to evaluate the ability of saRNAs to initiate an infectious cycle (i.e., infectivity) by quantifying the number of payload-expressing cells. To this end, BHK-21 cells were transfected with saRNAs carrying selected mutations and cells expressing GFP were counted in a FluoroSpot Analyzer the day after transfection (Figure 4). Similar to luciferase measurements, a significant reduction in the number of cells expressing GFP was observed for m^1^ -RNAs compared to unmodified RNAs, while incorporation of m^5^C had no apparent impact on counts. Altogether, our data suggest that incorporation of m^1^ negatively affected the self-amplifying capacity of saRNAs at the stage of RNA synthesis initiation.

**Figure 4.**
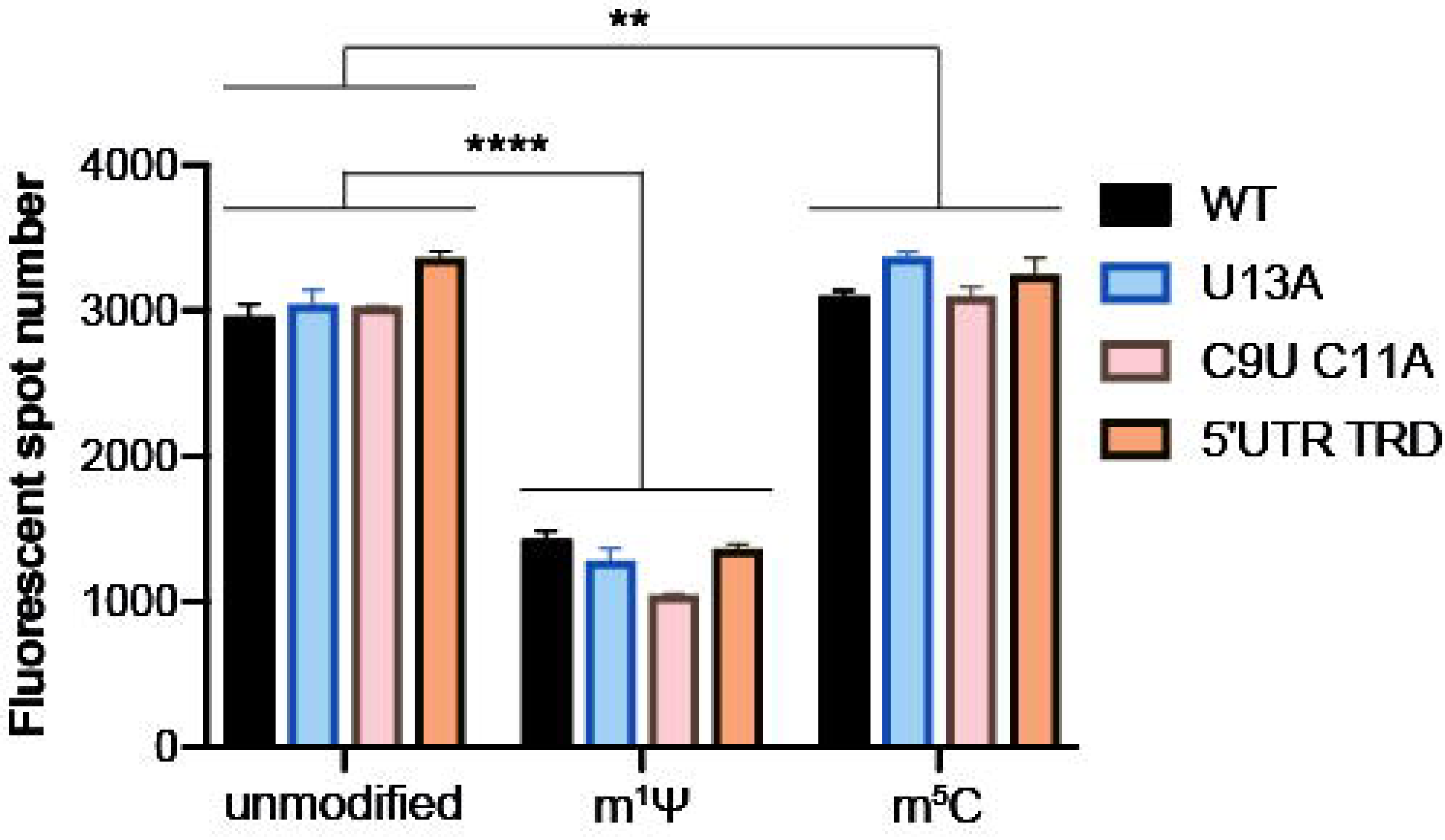
m^1^ψ modification impairs VEEV replicon infectivity. Bar graph showing foci counts in BHK-21 cells transfected with 80 ng of unmodified RNA or RNAs modified with m^1^ψ or m^5^C. GFP positive foci were numbered 24h post-transfection in an ImmunoSpot Analyzer and quantification was used as a measure of RNA infectivity. The main effect of modified nucleotides vs. unmodified was compared using Dunnett’s test. ns p> 0.05; **p ≤ 0.01; ****p ≤ 0.001. Error bars indicate the standard deviation.

The sets of 5’ and 3’UTR mutants were also tested in A549 cells (Figure S2). Similar to BHK-21 cells, the mutations did not affect Renilla luciferase activity 1 hour post-transfection nor firefly luciferase activity 24 hours post-transfection, compared to wild type modified and unmodified RNAs. The exception was the poly(U) deletion in the 3’UTR, which resulted in a drastic 10-to 50-fold reduction in firefly luciferase levels for all the RNAs analyzed.

In conclusion, saRNAs carrying mutations in the 5’ and 3’ UTR sequences of TC-83 VEEV retained the ability to replicate efficiently, however, none of the mutants was able to overcome the impact of m^1^ψ incorporation.

### Modulation of VEEV RNA-dependent-RNA-polymerase activity

Because the design of the UTR mutants was not suitable to overcome the impact of modified nucleotide incorporation on saRNA replication, we aimed at optimizing replicase function by modulating the RdRp activity of nsp4. It has been previously reported that uridine analogues are incorporated with lower fidelity than unmodified uridine during *in vitro* transcription [26]. In addition, synthetic nucleotide analogues used as anti-viral agents have been shown to be mutagenic [27]. Based on these observations, we speculated that the incorporation of modified analogues would increase the mutation rate due to their mutagenic effect during *in vitro* transcription of the input RNA and the first round of RNA synthesis by the viral RdRp. Hypermutated RNAs would drive the viral replicon beyond the threshold of viability explaining the impact of modified nucleotide incorporation on replicon RNA amplification. If that were the case, we reasoned that changes in the RdRp fidelity could aid in diminishing error rate during RNA synthesis. For different RNA viruses, growth in the presence of nucleotide analogues has led to the identification of residues in RdRp that are associated with polymerase fidelity [28,29]. For instance, mutations associated with anti-viral resistance to the nucleotide analogues ribavirin and favipiravir map to viral RNA polymerases. In the case of chikungunya virus (CHIKV), the change of Cys residue at position 483 to Tyr (C483Y) conferred resistance to ribavirin and was related to a high fidelity phenotype [15]. Interestingly, residue 483 appears to be a key modulator of polymerase fidelity, as screening of amino acid substitutions at this position also identified viral variants with a low fidelity phenotype. In addition, resistance of CHIKV to favipiravir arose from acquisition of Lys to Arg substitution at position 291 (K291R) of nsP4 [16].

To address the impaired replication rate of modified saRNA through modulation of polymerase fidelity, we introduced the mutations associated with CHIKV resistance to ribavirin and favipiravir into the analogous VEEV nsp4 residues (C482Y and K290R, respectively). In addition, we engineered a replicon containing the C482G substitution, associated with a low fidelity phenotype in CHIKV [30]. We used the reporter saRNA carrying Renilla and firefly luciferase genes to evaluate RNA translation and replication (Figure 5A). In addition, we engineered mutations into a new reporter VEEV saRNA consisting of a first ORF coding for the uninterrupted sequence of replicase proteins and a second ORF coding for GFP as a payload to evaluate RNA infectivity (Figure 5B). RNAs were *in vitro* transcribed with unmodified nucleotides or with m^1^ψ and transfected into BHK-21 cells. Renilla and firefly luciferase activities were measured at 1- and 24-hours post-transfection to assess RNA translation and replication, respectively (Figures 5A). One hour post-transfection, levels of Renilla luciferase activity were similar for the wild type (nsp4 WT) and the nsP4 mutant replicons, for both the unmodified and the m^1^ψ-RNAs, indicating that mutations in nsP4 did not affect translation (Figure 5A, top). As already noted, incorporation of m^1^ψ resulted in a moderate decrease Renilla luciferase activity. At 24 hours post-transfection, all the unmodified RNAs showed comparable levels of firefly luciferase activity (Figure 5A, bottom). In contrast, for m^1^ψ-RNAs, the C482Y and K290R mutants displayed a 2-fold increase in firefly luciferase levels, while the low fidelity C482G mutant displayed similar luciferase levels compared to the wild type RNA (Figure 5A, bottom). The number of foci paralleled luciferase activity (Figure 5C). The unmodified RNA of the high fidelity C482Y mutant displayed foci counts similar to the nsp4 WT RNA, while the m^1^ψ-RNA showed a 2-fold increase compared to the nsp4 WT m^1^ψ-RNA. Interestingly, the K290R mutant showed an improved ability to initiate a replicative cycle, yielding 0.5-fold and 10-fold increase in foci counts compared to the nsp4 WT for the unmodified and the m^1^ψ-RNAs, respectively. The fact that the positive effect of the mutation was superior for modified than for unmodified RNAs suggests that tuning of the polymerase function might be used as a strategy to address the impaired function of nucleotide-modified saRNAs.

**Figure 5.**
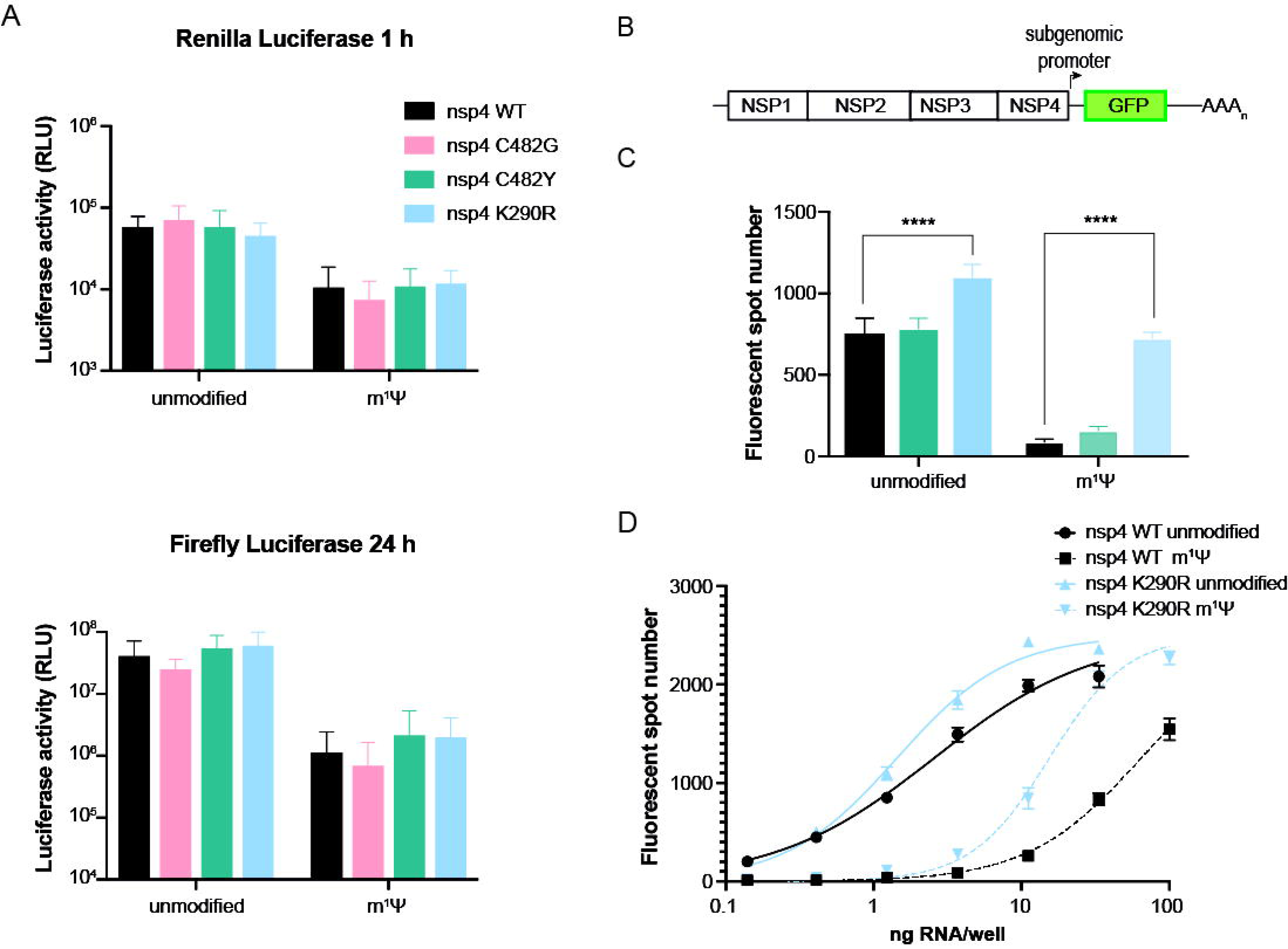
Mutations associated with increased RdRp fidelity partially rescue the impact of m^1^ψ modification on the self-amplifying ability and infectivity of VEEV replicon. (A) Bar graphs showing luciferase activity of BHK-21 cells transfected with 80 ng of unmodified RNA or m1ψ-modified RNAs of the wild type or nsp4 mutants C482G associated with a low fidelity phenotype, and C482Y and K290R associated with a high fidelity phenotype. Renilla luciferase activity was measured at 1h post-transfection (top) and firefly luciferase activity was measured at 24h post-transfection (bottom). (B) Schematic representation of a reporter VEEV TC-83 replicon expressing GFP. The GFP coding sequence was inserted in place of the viral structural proteins in the second ORF of the genome. (C) Bar graph showing foci counts in BHK-21 cells transfected with 1 ng of unmodified RNA or 10 ng of m1ψ modified RNA of the wild type and nsp4 mutants associated with a high fidelity phenotype. GFP positive foci were numbered 24h post-transfection in an ImmunoSpot Analyzer and quantification was used as a measure of RNA infectivity. Statistical analysis was performed using two-way ANOVA, followed by Tukey’s multiple comparisons test. ns p> 0.05; ****p ≤ 0.001. One representative experiment out of two is shown. Error bars indicate the standard deviation. (D) Dose-response curves of infectivity in BHK-21 cells transfected with increasing amounts of unmodified or m1ψ modified RNAs of the wild type or nsp4 K290R mutant VEEV-GFP replicon. GFP positive foci were numbered 24h post-transfection in an ImmunoSpot Analyzer. One representative experiment out of three is shown.

To further characterize the phenotype of the mutant K290R, we transfected increasing amounts of saRNA into BHK-21 cells (Figure 5D). After 24 hours, we counted the number of foci to estimate the 50% effective dose (ED50), defined as the amount of transfected saRNA necessary to reach 50% of the maximum foci number. For the unmodified nsp4 WT saRNA, the ED50 was 2.5 ng and the incorporation of m^1^ψ increased the ED50 about 25-fold (Table 1), confirming that m^1^ψ diminishes the number of GFP-expressing cells. The unmodified K290R mutant showed a 2-fold lower ED50 than the nsp4 WT counterpart, indicating that the mutation provides an advantage for heterologous protein expression. Remarkably, the incorporation of m^1^ into K290R saRNA showed a 10-fold increase in the ED50. Compared to the 25-fold increase observed for the nsp4 WT saRNA, this result suggests that the mutation partially counteracts the negative effect of the nucleotide analogue on protein expression. Remarkably, K290R mutant m^1^ψ-saRNA reached maximum foci counts at the plateau similar to the unmodified RNA, although at higher RNA doses. Thus, we were able to fully overcome the negative effect of nucleotide analogues by combining the K290R mutation with higher doses of saRNA. Altogether, mutation K290R in nsp4 alleviated the effect of m^1^ψ-incorporation, suggesting that modulation of RdRp activity may serve as an approach to extend the use of modified nucleotides to saRNA-based therapeutics.

**Table 1.** ED_50_ values estimated for WT and K290R mutant.

|  | WT | K290R |
| --- | --- | --- |
| unmodified | 2.554 | 1.435 |
| m1ψ | 64.16 | 15.38 |

## DISCUSSION

In this study, we investigated the effect of nucleotide analogues on saRNA payload expression. We found that m^1^ψ, ψ and m^5^C incorporation moderately increased RNA translation and stability. While m^1^ψ- and ψ negatively affected saRNA replication, saRNAs incorporating m^5^C showed similar replication levels than unmodified saRNAs. Impaired replication was not overcome by modifying the UTRs, likely due to the strict sequence and structure requirements for promoter activity. Importantly, the impact of m^1^ψ- and ψ on replication was partially reverted by point mutations that are associated to increased viral polymerase fidelity.

While incorporation of modified nucleotides is extensively used to improve translation efficiency, enhance stability, and reduce innate immune responses of mRNA vaccines, incorporation into saRNA appeared to be deleterious to payload expression in animal studies. In fact, expression of a human monoclonal antibody against ZIKV, driven from a TC-83 derived replicon, was undetectable in mice inoculated with the ψ-modified RNA formulated in lipid nanoparticle carrier [10]. Furthermore, induction of an immune response against SARS-CoV-2 was completely abrogated by the incorporation of m^1^ into the saRNA [11]. Here we gained insight into the molecular bases leading to the reported failure of modified saRNAs performance. Our findings pinpoint the impact of modified nucleotide incorporation to replication complex function. Strikingly, ψ- and m^1^ψ-modified RNAs showed >100-fold reduction at the stage of RNA replication compared to the unmodified RNA. Interestingly, m^5^C had no apparent effect on RNA replication. On the other hand, and in line with previous observations, incorporation of ψ, m^1^ψ, or m^5^C, resulted in a moderate increase of translation levels and stability of saRNAs, as evidenced after transfection of interferon competent A549 cells. Remarkably, the effect was observed only in the context of the saRNA with impaired RdRp activity. As we were able to detect saRNA replication as early as 2 hours after transfection, we interpreted that the negative impact of nucleotide incorporation at the stage of RNA synthesis masked the stimulatory effect on translation and stability.

Previous research has explored the use of m^5^C as an alternative modification to m1ψ. m^5^C modification has been found to enhance RNA translation efficiency [5] and reduce innate immune responses [4,17]. Immunogenicity studies have demonstrated that m^5^C-modified RNAs display decreased activation of human dendritic cells and cells stably expressing TLR3, TLR7, or TLR8 [4]. Our data show that, unlike uridine analogs, m^5^C did not impact the self-amplifying ability of a replicon RNA, and together with the previous immunogenicity studies, highlight the potential of m^5^C-modified saRNA as a viable approach for vaccine applications.

Besides the immunomodulatory effects of modified nucleotide incorporation into RNAs, the replacement of specific uridines with ψ can influence the structural stability. Seminal observations indicated that replacement of uridines by ψ stabilized the conformation of an RNA oligonucleotide [31]. Later studies pointed out that incorporation of ψ altered the function of structured RNA regulatory elements, such as IRESs[18]. Recently, a systematic analysis where specific uridines were replaced by ψ, in the context of a functional riboswitch, suggested that the impact of substituting particular uridines with pseudouridines is highly contingent upon its precise location, resulting in effects that can range from destabilizing to stabilizing [19]. Structured RNA elements are crucial to the self-amplifying ability of viral replicons as they can function controlling translation and immune sensing and as promoters for RNA synthesis. Our approach of replacing specific uridines and cytidines to avoid modified nucleotides incorporation in the 5’SL of the TC-83 replicon did not alleviate the impaired function of modified RNAs. Given the complex organization of virus RNA genomes displaying overlapping signals that can function in alternative RNA conformations, we cannot rule out that incorporation of modified nucleotides had no effect on the dynamics of these structures.

Modified nucleotides can have mutagenic potential arising from the base-pairing patterns and altered interactions between modified nucleotides and RNA polymerases during transcription [26,32]. For instance, increased error rates were reported in the *in vitro* synthesis of RNAs incorporating ψ by T7 RNA polymerase and to a lesser extent in RNAs incorporating m1ψ. Also, an increase in the error rate of first strand RNA synthesis was observed for reverse transcriptases using modified RNAs as template [26]. The saRNA vaccine technology relies on the *in vitro* transcription of an RNA molecule that is used as the template for RNA synthesis by the virus replicase upon delivery into a target cell. Therefore, *in vitro* transcription and complementary RNA synthesis can have a cumulative effect in the error rate impairing the self-amplifying ability of a modified saRNA that is pushed to error catastrophe. Here, we found that mutations in the nsp4 RdRp of VEEV TC-83 that were originally selected by passaging the closely related chikungunya virus in the presence of nucleotide analogs as mutagens and are linked to a high fidelity phenotype were able to partially overcome the impact of uridine analog incorporation on RNA replication. Our results suggest that nucleotide modified RNA templates increase the error rate of the viral replicase. However, research assessing nucleotide misincorporation by RdRps copying modified RNAs to synthesize a complementary strand is still needed.

Overall, together with previous studies, our data encourage the incorporation of m^5^C in saRNA vaccine candidates as they display appropriate balance of the immune response elicited by the RNA molecule and it has a minor impact in the self-amplifying ability of virus replicons. Furthermore, we uncovered a previously unappreciated link between polymerase fidelity and replication of m^1^ψ-modified self-amplifying RNA molecules. Therefore, engineering of the replicase function may serve as a strategy to improve the potency of saRNA platforms and improve efficacies in future clinical studies.

## MATERIALS AND METHODS

### Cells

Cell lines were grown at 37 °C in a 5% CO_2_ atmosphere. BHK-21 cells (ATCC, CCL-10) were grown in alpha-modified Eagle’s minimum essential medium (α-MEM) supplemented with 10% fetal bovine serum (FBS) and penicillin−streptomycin antibiotics. Human lung cell line A549 (ATCC CL-185) was cultured with DMEM supplemented with 10% FBS and antibiotics.

### Constructs

The VEEV-GFP replicon was derived from a plasmid encoding the VEEV TC-83 sequence under the control of a T7 promoter by inserting GFP coding sequence downstream of the subgenomic promoter and upstream of the 3’UTR. The VEEV replicon with the luciferase reporters was constructed by restriction free cloning of Renilla luciferase coding gene in the hypervariable region of nsP3 (after the codon 374), and firefly luciferase coding gene downstream of the subgenomic promoter and upstream of GFP coding gene in VEEV-GFP. 5’UTR, 3‘UTR and nsp4 mutants were generated using a restriction-free approach [33] involving: (i) a first PCR to amplify the insert sequence with designed mutations (Supplementary Table 1); (ii) a second PCR using the insert sequence as a mega-primer and VEEV replicon plasmid as a template, and (iii) DpnI digestion and transformation of the second PCR product.

### In vitro transcription and purification of saRNAs

The VEEV replicon reporter was linearized by restriction digestion with BspQI at the precise 3’ end of SAM sequences and purified by a EasyPure PCR Purification Kit (TransGen Biotech). Linearized DNAs were used as templates for *in vitro* transcription of capped RNAs with HiScribe™ T7 High Yield RNA Synthesis Kit (New England Biolabs) in the presence of CleanCap® Reagent AU (TriLink Biotechnologies). For modified RNAs, UTP was fully substituted with either pseudouridine-5’-triphosphate (TriLink Biotechnologies) or N1-Methylpseudouridine-5’-Triphosphate (TriLink Biotechnologies), or CTP was fully substituted with 5-methylcytidine-5’-triphosphate (TriLink Biotechnologies) in the *in vitro* transcription reactions. The reaction mixes were incubated at 37 °C for 2 h. Then, the DNA template was digested with DNase (Invitrogen) at 37 °C for 30 min and the RNA transcripts were precipitated with lithium chloride overnight at -70°C. Next, RNAs were recovered by centrifugation at 12000 xg for 1 hour at 4°C, washed with 70% ethanol and centrifugation at 12000 g for 5 minutes at 4°C, and resuspended in nuclease-free water. The quality of the RNA samples was analyzed by 1% agarose gel electrophoresis. The concentration of RNA was determined in a Nanodrop spectrophotometer (Thermo Fisher Scientific).

For capping with a dinucleotide cap analog, in vitro transcription was performed with a NTP mix consisting of CTP, GTP, and UTP, and ATP and ^m7^GpppA_m_ cap analog (New England Biolabs) at a 1:5 ratio.

### RNA transfection, measurements of firefly and Renilla Luciferase activities, and counting of GFP-expressing foci

BHK or A549 cells were seeded into 96-well plates at 3.5 × 10^4^ cells per well. After 24 hours, *in vitro* transcribed saRNAs were transfected into cultured cells using 0.2 μl of Lipofectamine-2000 (Invitrogen) according to the manufacturer’s instructions. The precise amount of RNA used in each experiment is indicated in the corresponding figure legend. Firefly and Renilla Luciferase activities were sequentially measured in a GloMax luminometer (Promega) using the Dual-Luciferase Reporter Assay System (Promega). Briefly, transfected cells were lysed in 100 μl of 1X Passive Lysis Buffer (Promega), and the firefly luciferase reporter was measured first by adding Luciferase Assay Reagent II (LAR II) to the well. After quantifying firefly luciferase activity, Stop & Glo Reagent was added to simultaneously quench the firefly reaction and initiate the Renilla luciferase reaction.

To count GFP-expressing foci, BHK and A549 cells were transfected as above and at 24 hours post-transfection, the culture media was removed and 30 µL of 1X PBS was added to the wells. The fluorescent spots were quantified using an ImmunoSpot Analyzer (Cellular Technology).

## Data Analysis

Statistical analysis was performed using GraphPad Prism software (version 9.2).

## Acknowledgements

DA and CVF are members of CONICET. As Senior Director working for GreenLight Biosciences MS proposed and executed a research sponsor agreement between GLB and UNSAM.

---

**SUPPLEMENTARY TABLE 1.**
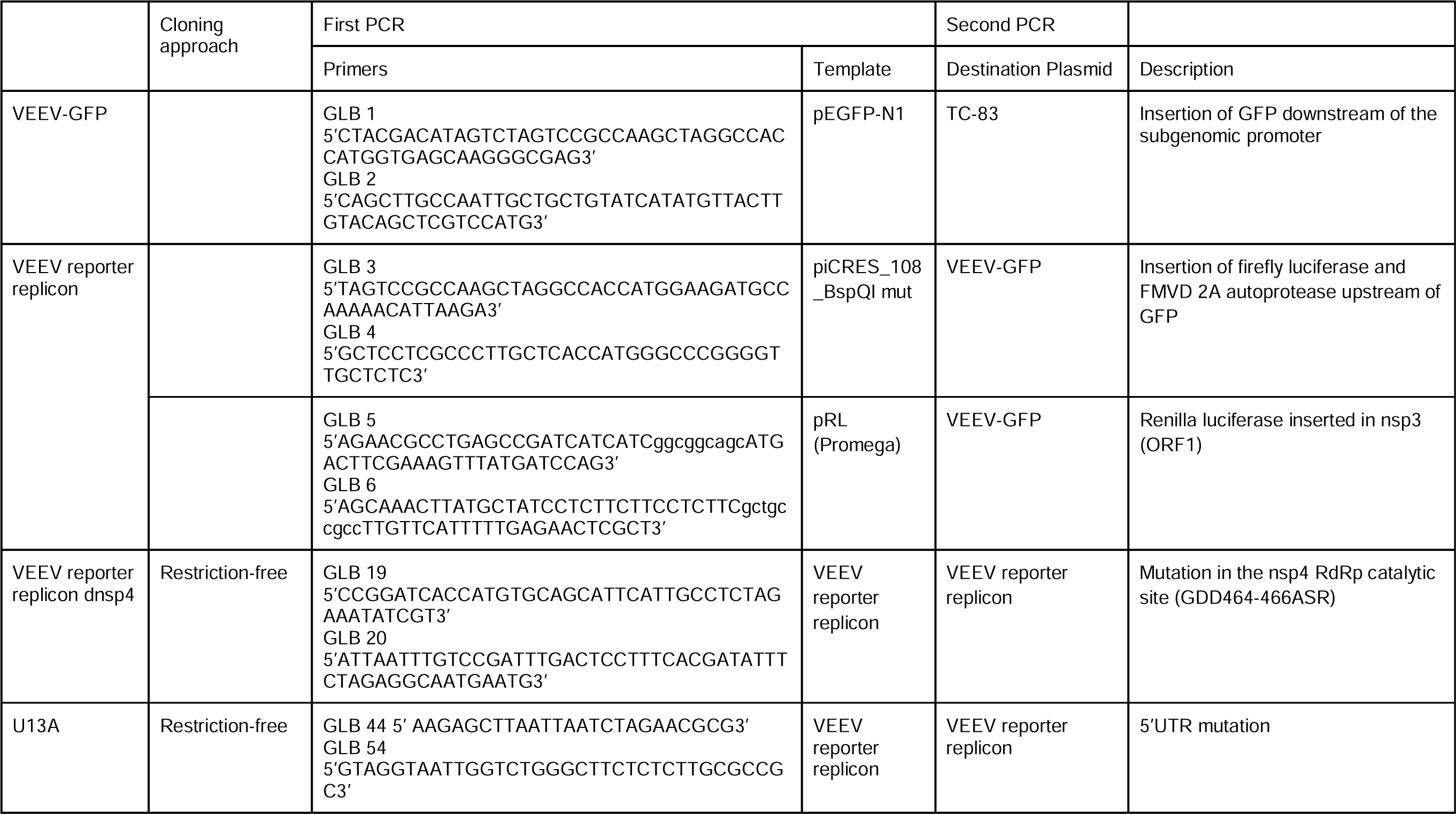
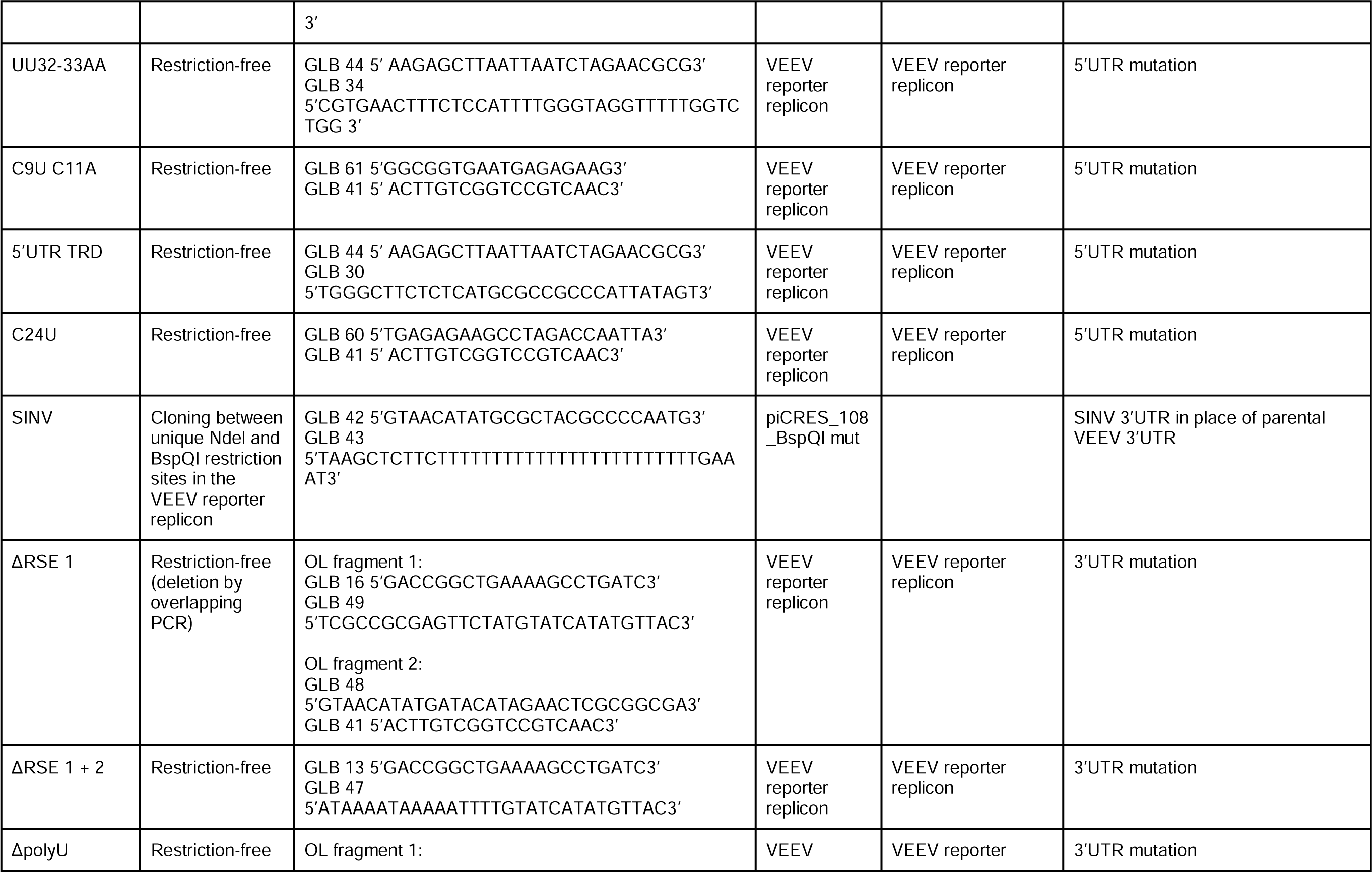
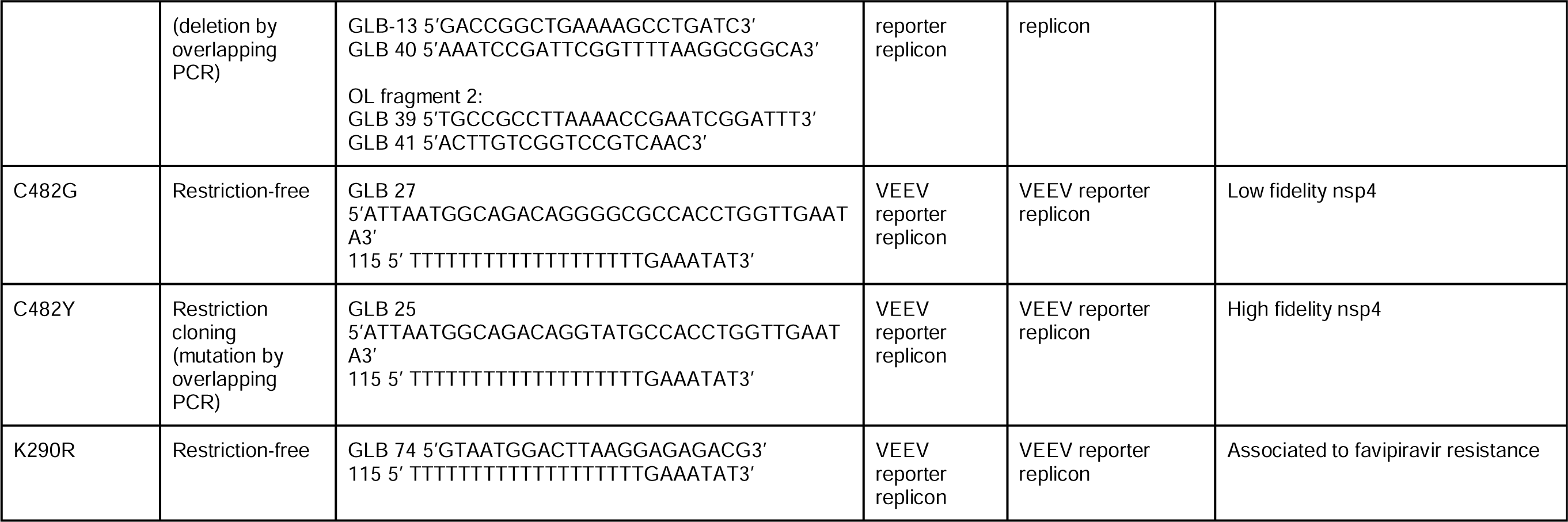

## FIGURE LEGENDS

**Figure S1. Effect of nucleotide analogues on VEEV reporter translation and replication in A549 cells.** Bar graph displaying Renilla luciferase activity of dnsp4 mutant (A) and firelfy luciferase activity for WT (B) at 2-, 8- and 24-hours post-transfection of A549 cells with VEEV replicon RNA transcribed in vitro with unmodified nucleotides or with ψ, m1ψ or 5mC. Two independent experiments with two replicates per condition were analyzed. Statistical analysis was performed using two-way ANOVA, followed by Dunnett’s test to compare the main effect of modified nucleotides vs. unmodified. ns p> 0.05; *p ≤ 0.05. Bars indicate the standard deviation.

**Figure S2. Effect of 5’UTR and 3’UTR mutants on VEEV replicon replication in A549 cells.**

Bar graphs showing luciferase activity of A549 cells transfected with 80 ng of unmodified RNA, m1ψ- or 5mC-modified RNAs. Renilla luciferase activity was measured at 1h post-transfection (left) and firefly luciferase activity was measured at 24h post-transfection (right). Statistical analysis was performed using two-way ANOVA, followed by Dunnett’s multiple comparisons test. Lines compare the main effect of modified nucleotides against unmodified. ns p> 0.05; *p ≤ 0.05; **p ≤ 0.01; ***p ≤ 0.001; ****p ≤ 0.001. Error bars indicate the standard deviation.

