## Supplementary figures and images for "Improvement of the potency of a N1-methylpseudouridine-modified self-amplifying RNA through mutations in the RNA-dependent-RNA-polymerase"

### Supplemental Figure 1

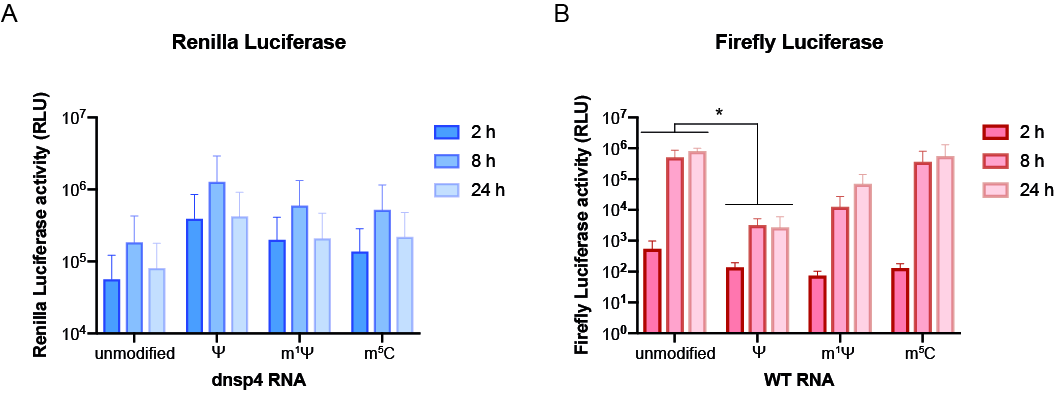

### Supplemental Figure 2

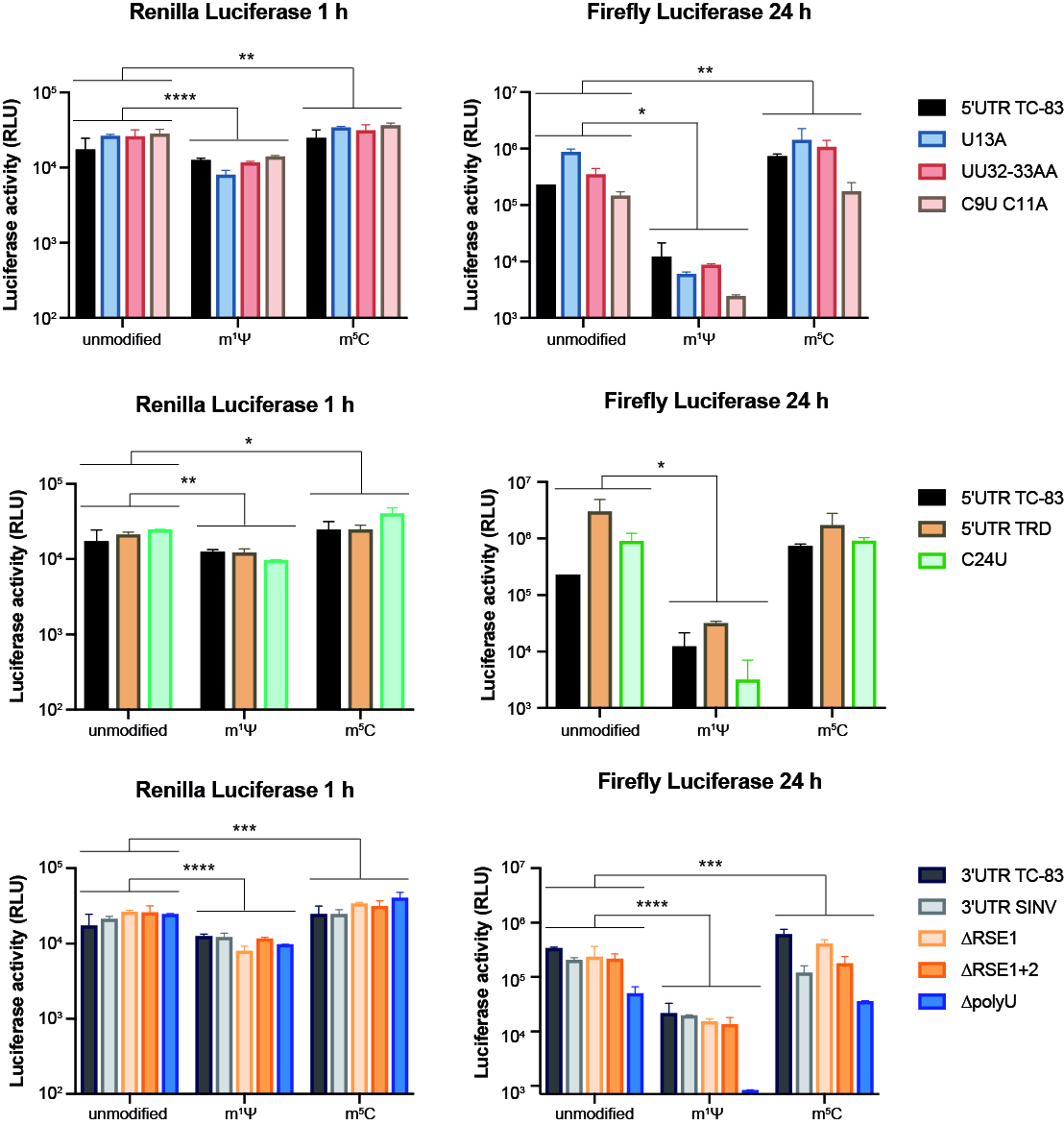
